# Expressing the pro-apoptotic Reaper protein via insertion into the structural open reading frame of Sindbis virus reduces the ability to infect *Aedes aegypti* mosquitoes

**DOI:** 10.1101/2022.01.13.476239

**Authors:** Alexis Carpenter, Rollie J. Clem

## Abstract

Arboviruses continue to threaten a significant portion of the human population, and a better understanding is needed of the determinants of successful arbovirus infection of arthropod vectors. Avoiding apoptosis has been shown to be one such determinant. Previous work showed that a Sindbis virus (SINV) construct called MRE/rpr that expresses the pro-apoptotic protein Reaper via a duplicated subgenomic promoter had a reduced ability to orally infect *Aedes aegypti* mosquitoes at 3 days post-blood meal (PBM), but this difference diminished over time as virus variants containing deletions in the inserted *reaper* gene rapidly predominated. The goal of this study was to generate a SINV construct that more stably expressed Reaper, in order to further clarify the effect of midgut apoptosis on disseminated infection in *Ae. aegypti*. We did this by inserting *reaper* as an in-frame fusion into the structural open reading frame (ORF) of SINV. This construct, MRE/rprORF, successfully expressed Reaper, replicated similarly to MRE/rpr in cell lines, and induced apoptosis in cultured cells and in mosquito midgut tissue. Mosquitoes that fed on blood containing MRE/rprORF developed less midgut and disseminated infection when compared to MRE/rpr or a control virus up to at least 7 days PBM, when less than 50% of mosquitoes that ingested MRE/rprORF had detectable disseminated infection, compared with around 80% or more of mosquitoes fed with MRE/rpr or control virus. However, virus titer in mosquitoes infected with MRE/rprORF was not significantly different from control virus, suggesting that induction of apoptosis by expression of Reaper by this method can reduce infection prevalence, but if infection is established then apoptosis induced by this method has limited ability to continue to suppress replication.

## INTRODUCTION

Recent decades have seen the emergence and reemergence of a number of significant arboviral diseases such as dengue, Zika, West Nile, yellow fever and chikungunya (1–3). Arboviruses, which are transmitted through the bite of an infected arthropod vector such as a tick or mosquito, are expected to become more significant in the future and it is predicted that climate change will increase incidence of disease by impacting vector geographical range, feeding behavior and survival (4–6). New ways of protection from these diseases are needed as vaccines are not available for many of these diseases and there is increasing insecticide resistance in some vectors (7–9).

An alternative to traditional means of controlling arboviral diseases is to prevent productive infection of the vector. Tissue barriers in the vector such as the midgut present an obstacle for viruses to overcome and inhibiting escape from midgut tissue would prevent disseminated infection and thus the spread of these viruses (10). Several pathways and processes may be considered which when altered could prevent disseminated infection, and improved knowledge of these pathways could lead to new strategies of vector infection control. One cellular process which shows potential promise in preventing disseminated infection is apoptosis. Apoptosis, a specific type of programmed cell death, has been shown to be an important antiviral pathway in insects (11,12). The insect apoptosis pathway has been best studied in *Drosophila melanogaster*, but a similar pathway has been demonstrated in the disease vector *Aedes aegypti* (13–16). During apoptosis, activated initiator caspases cleave and activate effector caspases, which are responsible for cleaving target proteins in the cell, leading to death. Activation of initiator caspases is prevented by inhibitor of apoptosis (IAP) proteins, the action of which can be overcome by IAP antagonists such as Reaper. The result is a carefully controlled mechanism which prevents unnecessary cell death but promotes cell death in response to activating stimuli such as viral infection.

The role of apoptosis in protecting insects against viral infections brings up the question of what role it plays in vector competence for arboviruses. Several studies have implicated apoptosis as a significant factor in preventing viral escape from the midgut. For example, enhanced midgut apoptosis has been associated with a *Culex pipiens pipiens* strain of mosquitoes that were refractory to West Nile virus infection (17). Additionally, a SINV construct that expressed the *Drosophila* IAP antagonist Reaper was shown to less effectively infect and disseminate from the *Aedes aegypti* midgut, although viruses with deletions in the *reaper* insert rapidly predominated (18). Consistent with these results, one study showed that some proapoptotic genes were more highly expressed in a refractory strain compared to a susceptible strain of *Aedes aegypti* (19) while another study found increased rapid induction of apoptosis in mosquitoes that were less susceptible to dengue virus serotype 2 compared to a more susceptible strain (20). However, it has also been suggested that apoptosis may weaken the barrier that the midgut provides and thus allow viruses to pass through more easily. One study found that knocking down AeIAP1 expression in *Aedes aegypti* and then feeding them with SINV led to increased disseminated infection (21). Due to a high rate of mosquito mortality in these AeIAP1 knockdown mosquitoes, it was hypothesized that the level of apoptosis was drastically increased, greatly reducing the structural integrity of the midgut. It is possible that some level of apoptosis is critical to preventing virus passage through tissues but if apoptosis levels are too high, viral spread is promoted through gaps in the structure.

A previous series of studies by our group aimed to study how inserting the *Drosophila* pro-apoptotic gene *reaper* into SINV and then infecting *Aedes aegypti* would affect rates of disseminated infection (18,22). The construct used in these earlier studies, called MRE/rpr, expressed Reaper via a duplicated subgenomic promoter located between the nonstructural and structural genes. The MRE/rpr virus strongly induced apoptosis and had decreased virus yield compared to control viruses in cultured mosquito cells (22). When mosquitoes were fed MRE/rpr, it was found that there was less disseminated infection compared to control virus at early timepoints, but by 7 days post-blood meal (PBM) there was no significant difference between MRE/rpr and control. The reason for this was found to be that a significant proportion of the MRE/rpr population by 7 days PBM had deletions in the *reaper* insert, rendering it non-functional (18). This result indicated strong negative selection against Reaper expression. Indeed, even at early timepoints over half of the mosquitoes developed disseminated infection, which may have been due to the strong selective advantage of mutants lacking functional *reaper*.

To provide more stable expression of Reaper protein, we decided to insert the *reaper* gene into the structural ORF of SINV, a method which has been previously shown to allow more stable insertions into the SINV genome (23,24). By inserting the sequence into the structural ORF between the capsid gene and PE2, as opposed to the duplicated subgenomic promoter region, we hoped to increase selective pressure for retaining the insert because deletions in *reaper* would be more likely to negatively impact the correct expression of critical viral structural proteins. To allow proper synthesis of the viral structural proteins, we utilized the autoproteolytic function of the SINV capsid protein and the ribosomal skipping function of foot and mouth disease virus (FMDV) 2A (25) to cotranslationally release the Reaper protein. Additionally, we employed a ubiquitin fusion strategy which has been employed successfully to generate proteins with precise N-terminal sequences (26,27). This ensured that we would not impact the N-terminus of Reaper, which has been shown to be critical for binding IAPs (28).

Using this SINV construct that more stably expressed Reaper, we then re-examined the effect of Reaper expression on establishment of midgut infection and dissemination from the midgut. Our results provide deeper insights into the effects of apoptosis on SINV infection in *Ae. aegypti*.

## METHODS

### Cells

BHK-21 cells were maintained at 37°C with 5% CO_2_ in Dulbecco modified Eagle medium (DMEM, Gibco) plus 10% fetal bovine serum (FBS, Atlanta Biologicals). C6/36 cells were maintained at 27°C in Liebovitz’s medium (Gibco) plus 10% FBS.

### Insect rearing

Orlando strain *Aedes aegypti* mosquitoes (obtained in 2008 from James Becnel, USDA ARS, Gainesville, FL) were reared in a 27°C incubator with 80% humidity and a 12-hr light-dark cycle. To obtain eggs used in experiments, females were allowed to feed on defibrinated sheep’s blood (Colorado Serum Company) using a Hemotek feeding system (Hemotek Ltd.). Adult mosquitoes were maintained on raisins and water.

### Plasmid design and construction

A previously described plasmid containing a fragment of 5’dsMRE16ic extending from the NotI site to the AvrII site as well as microRNA target sites and a 2A self-cleaving peptide sequence ligated into a pGEM-T backbone was used as the starting plasmid (24). This plasmid was digested with BstEII and AfIII and the intervening sequence containing the miRNA target sites was replaced with the following sequence (synthesized by Genewiz) containing ubiquitin (human K48R), the *Drosophila reaper* sequence (lacking the initiator methionine) and an HA tag: (TGGAATAGCAAGGGAAAGACCATCAAGACGACGCCCGAAGGGACAGAGGAATGGTCAGCAGCACTCGAGATGCAGATCTTCGTCAAGACGTTAACCGGTAAAACCATAACTCTAGAAGTTGAACCATCCGATACCATCGAAAACGTTAAGGCTAAAATTCAAGACAAGGAAGGCATTCCACCTGATCAACAAAGATTGATCTTTGCCGGTAGGCAGCTTGAGGACGGTAGAACGCTGTCTGATTACAACATTCAGAAGGAGTCCACCCTGCACCTGGTCCTCCGTCTCAGAGGTGGTGCAGTGGCATTCTACATACCCGATCAGGCGACTCTGTTGCGGGAGGCGGAGCAGAAGGAGCAGCAGATCCTTCGCTTGCGGGAGTCACAGTGGAGATTCCTGGCCACCGTCGTCCTGGAAACCCTGCGCCAGTACACTTCATGTCATCCGAAGACCGGAAGAAAGTCCGGCAAATATCGCAAGCCATCGCAATACCCATACGATGTTCCAGATTACGCTGGATCCCAGCTGTTGAATTTTGACCTT). A control plasmid was generated in the same way, but the intervening sequence was replaced with a synthesized sequence not containing *Drosophila reaper* but still containing ubiquitin and an HA tag: (TGGAATAGCAAGGGAAAGACCATCAAGACGACGCCCGAAGGGACAGAGGAATGGTCAGCAGCACTCGAGATGCAGATCTTCGTCAAGACGTTAACCGGTAAAACCATAACTCTAGAAGTTGAACCATCCGATACCATCGAAAACGTTAAGGCTAAAATTCAAGACAAGGAAGGCATTCCACCTGATCAACAAAGATTGATCTTTGCCGGTAGGCAGCTTGAGGACGGTAGAACGCTGTCTGATTACAACATTCAGAAGGAGTCCACCCTGCACCTGGTCCTCCGTCTCAGAGGTGGTTATCCATACGATGTTCCAGATTACGCTGGATCCCAGCTGT TGAATTTTGACCTT). Both plasmids were then digested with NotI and AvrII and the purified fragments were ligated into 5’dsMRE16ic which had been digested with the same restriction enzymes. Proper insertion was verified by Sanger sequencing. The generation of MRE/rpr has previously been described (22).

### Virus production

Infectious clone plasmids were linearized using AscI (New England Biolabs) and then in vitro transcribed using the MEGAscript SP6 transcription kit (Thermo Fisher Scientific) with added cap analog (New England Biolabs). Following transcription, RNA was transfected into BHK-21 cells using Lipofectamine 3000 (Thermo Fisher Scientific). After two days, the media was removed and used to infect a T75 flask of C6/36 cells. After 5 days the resulting P2 virus stock was frozen in aliquots and titer was determined using TCID_50_ assay.

### TCID_50_ assay

BHK-21 cells were plated at a density of 1×10^4^ cells per well in a 96-well tissue culture plate in 100 µl of DMEM plus 10% FBS and supplemented with 15 µg per ml of penicillin/streptomycin (Invitrogen). Mosquito and cell samples were removed from -80° C and thawed on ice. Samples were then spun to remove debris and DMEM was used to make serial dilutions of each sample. Each dilution was transferred to five wells containing BHK-21 cells. After 5 days, each well was scored for cytopathic effects. The number of wells of each dilution scored as positive was used to determine TCID_50_ ml^-1^ and this was converted to PFU ml^-1^ by multiplying by 0.69 (29).

### Growth curves

For growth curves in C6/36 cells, the cells were plated at a density of 1×10^6^ cells per well in a 6-well plate in 2 ml of Leibovitz’s medium containing 10% FBS. For growth curves in BHK-21 cells, the cells were plated at a density of 5×10^5^ cells per well in a 6-well plate in 2 ml DMEM containing 10% FBS. Cells were allowed to recover for 2 hrs and then were infected with MRE/rprORF, MRE/rpr or MRE/control at a multiplicity of infection (MOI) of 0.1. The cells were placed on a rocker and the virus was allowed to adsorb for 1 hr. The media was then removed, and the cells were rinsed before replacing the media. For the cumulative growth curves, 100 μ l of media were sampled for analysis at 1, 2, 3 and 4 dpi. For the non-cumulative growth curves, 100 μ l of media were sampled at each time point for analysis and then the remaining media was removed, the cells were rinsed twice and then 2 ml of media were added. All samples were frozen and stored at -80°C until analysis by TCID_50_ assay.

### DNA fragmentation assay

C6/36 cells were plated at a density of 2 × 10^6^ cells per well in a 6-well plate in 2 ml Leibovitz’s medium containing 10% FBS. Cells were allowed to recover for 2 hrs and then were infected with MRE/rprORF, MRE/control, or 5’dsMRE16ic at an MOI of 1. After 48 hrs cells were removed from the 6-well plates and pelleted at 500 x g. They were washed twice with phosphate buffered saline (140 mM NaCl, 2.7 mM KCl, 10 mM Na2HPO4, 1.8 mM KH2PO4) and then resuspended in 200 μ l lysis buffer (0.1 M SDS, 0.1 M Tris pH 8.0, 0.05 M EDTA pH 8.0, 200 mg ml^-1^ Proteinase K) and incubated at room temperature for 1 hr. The samples were then extracted twice with phenol/chloroform and then ethanol precipitated. The pelleted DNA was resuspended in 100 μ l Tris-EDTA buffer (10 mM Tris-Cl, 1 mM EDTA) containing 100 μ g ml^-1^ RNase A (Thermo Fisher Scientific) and incubated at room temperature for 5 min. Twenty μ l of each sample was loaded into a 1.2% agarose gel containing 0.5 μ g ml^-1^ ethidium bromide. The gel was visualized using an AlphaImager gel imaging system. Sizes of the bands were compared to a Versaladder DNA ladder (Gold Bio).

### Mosquito infection for TCID_50_ and caspase assay

Prior to blood feeding, mosquitoes were placed in cups containing 20-30 females and 10% males and were only provided water for 24 hrs. MRE/rprORF, MRE/rpr and MRE/control stocks were diluted to 1.45 × 10^7^ PFU ml^-1^ with Liebovitz’s medium. The diluted virus stocks were then mixed 1:1 with defibrinated sheep blood and the mosquitoes were then allowed to feed on one of the virus blood mixtures using the Hemotek feeding system for 90 min. Fully engorged females were separated from unfed and partially fed females and males and were given water and raisins to feed *ad libitum*. At 3, 5, and 7 days PBM, the mosquitoes were cold anesthetized, and the midguts were dissected from the rest of the mosquito (the carcass). The midguts and carcasses used for TCID_50_ assays were placed in 1.5 ml tubes containing 200 μ l DMEM media containing 10% FBS and homogenized using disposable pestles. The samples were then frozen at -80° C. The midguts used for caspase assays were collected in 30 μ l of caspase reaction buffer (20 mM Hepes-KOH, pH 7.5, 50 mM KCl, 1.5 mMMgCl2, 1 mM EDTA, 1 mM EGTA, 1 mM DTT), homogenized with disposable pestles and stored at -80°C.

### Immunoblotting

C6/36 cells were plated at a density of 2×10^6^ cells per well in a 6-well plate containing 2 ml Leibovitz’s media plus 10% FBS. They were then infected with MRE/rprORF, MRE/control or 5’dsMRE16ic at an MOI of 10. After 24 or 48 hrs, the cells were rinsed three times with cold PBS. The plate was placed on ice and 100 ul of cold Laemmli sample buffer (Bio-Rad) was added to each well. The wells were scraped, and the lysate was collected in a 1.5 ml tube. The samples were then heated to 100°C for 5 min, centrifuged at 4°C and the supernatant was transferred to a new tube. SDS-PAGE was performed using 30 μ l of sample with 4-20% Bis-Tris gels (Genscript) in Tris-MOPS-SDS running buffer (Genscript), and proteins were transferred to PVDF membrane. After blocking for 1 hr in 5% dried skim milk in TBST, a 1:1000 dilution of the anti-HA (Biolegend) or anti-β-actin (Santa Cruz Biotechnology) primary antibody was incubated with the membrane overnight at 4°C with constant agitation. After incubation, the membrane was washed three times with TBST. A 1:15000 dilution of goat anti-mouse IgG-horseradish peroxidase secondary antibody (Thermo Fisher Scientific) was added and was incubated for 1 hr at room temperature with agitation. Bands were detected using SuperSignal West Pico PLUS Chemiluminescent Substrate (Thermo Fisher Scientific) and visualized using a LI-COR Western Blot Imager.

### Caspase assay

In the cell experiments, C6/36 cells were plated at a density of 2×10^6^ cells per well in a 6-well plate in 2 ml Leibovitz’s media plus 10% FBS. The cells were infected with MRE/rprORF, MRE/rpr or MRE/control at an MOI of 1. At 24 and 48 hpi the cells were rinsed with PBS and then collected in 400 µl of caspase reaction buffer. Mosquito midgut samples were collected in pools of 8 in caspase reaction buffer and the tissues were disrupted by sonication. The tubes were centrifuged, and the supernatant was moved to a new tube. All samples were frozen at -80°C until testing. Bradford protein assay (Bio-Rad) was used to determine protein concentration, and cell and mosquito samples were diluted to 100 µg ml^-1^. 50 µl of each sample were added to white 96-well plates (Costar) and incubated at 37°C for 15 min. Then 10 µl of Ac-DEVD-AFC (ApexBio) was added to each well at a final concentration of 20 µM. Cleavage of this fluorogenic substrate was monitored at an excitation wavelength of 405 nm and an emission wavelength of 535 nm using a Victor3 1420 Multilabel Counter (Perkins-Elmer). Readings were taken at 15 min intervals for a period of 1 hr.

## RESULTS

### MRE/rprORF virus construction and Reaper expression

To generate a SINV construct with enhanced stability of the pro-apoptotic gene *reaper*, we inserted this gene into the structural ORF of the 5’dsMRE16ic infectious clone (Fig. 1A) (30). The resulting construct, which we named MRE/rprORF, has *reaper* and flanking sequences inserted immediately after the capsid gene (Fig 1B). To ensure proper processing of both Reaper and the SINV capsid and E2 proteins, the first sequence inserted after the capsid gene was the first three codons of PE3, which are required for an intact capsid autoproteolytic cleavage site. However, this created a problem because these three amino acids would be fused to the Reaper protein, and the N-terminus of Reaper has been shown to be critical for its function in binding IAPs (28). Following normal translation of cellular Reaper, methionyl peptidases remove the N-terminal methionine, revealing an alanine that is required for binding to IAPs (31). To avoid this problem, we used a ubiquitin-Reaper fusion strategy. After the three PE3 codons, we inserted a *ubiquitin* gene immediately followed by the *reaper* gene (lacking the N-terminal methionine) with a hemagglutinin (HA) epitope tag at its C-terminus to facilitate detection of Reaper expression. Cellular proteases cleave ubiquitin fusion proteins immediately following the ubiquitin sequence (26). Thus the insertion of ubiquitin would result in expression of Reaper with amino-terminal alanine. Following *reaper*, we inserted the FMDV 2A sequence which causes ribosomal skipping (25) and thus should free the Reaper protein from the rest of the polypeptide chain. The result was a predicted Reaper-HA fusion protein expressed from the structural ORF that allows for intact viral structural protein processing. As a control virus we constructed MRE/control, which contained the same elements described above but lacked the *reaper* sequence (Fig. 1B). The other constructs used in this study were MRE/rpr (Fig. 1C), which has been previously described and expresses Reaper via a duplicated subgenomic promoter, and 5’dsMRE16ic, which also has been previously described and did not contain any of the inserted sequences other than the duplicated subgenomic promoter present in all of the constructs (18,22,30).

**Fig. 1:**
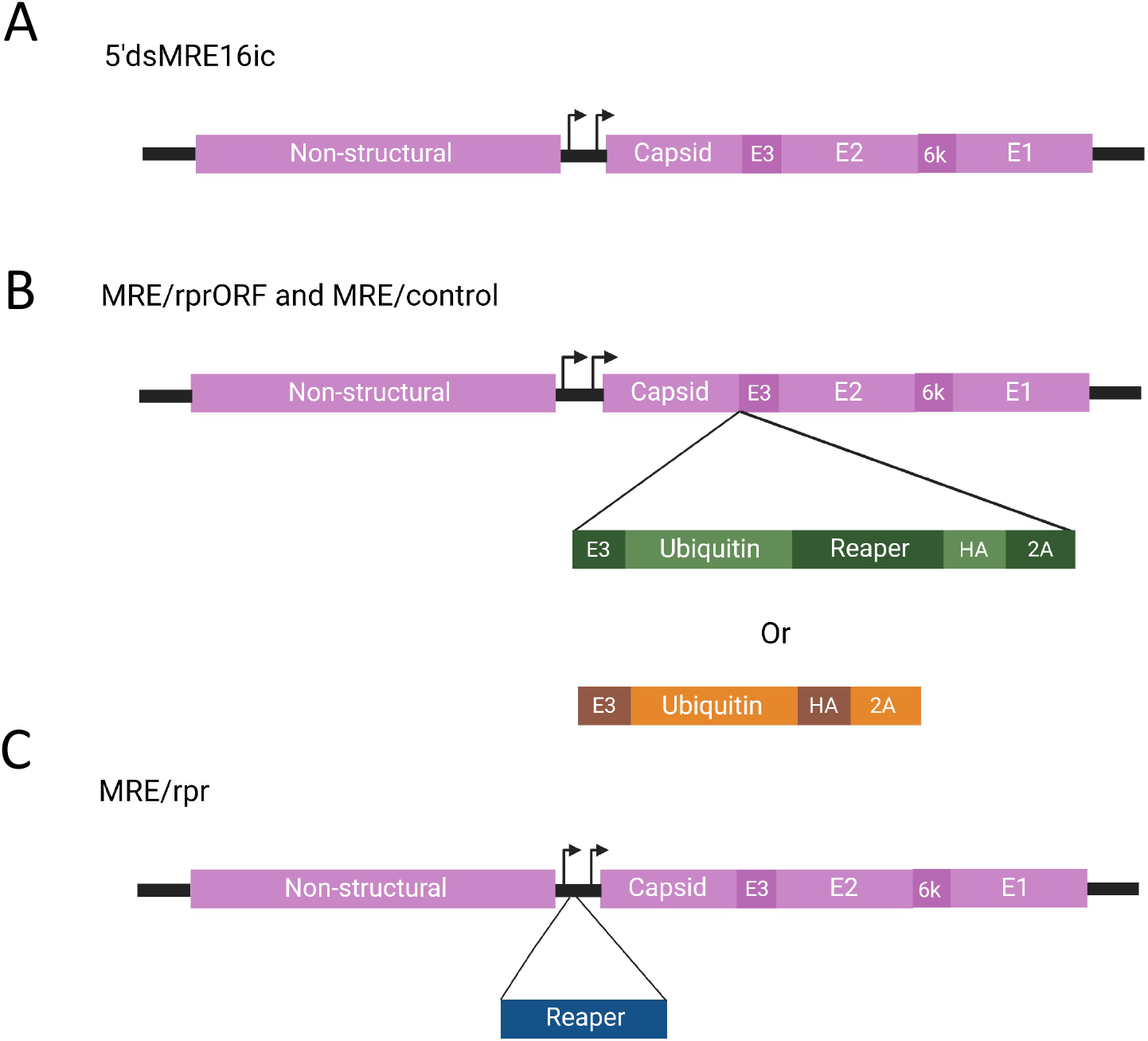
Diagrams of virus constructs used in this study. (A) 5’dsMRE16ic was used as the starting backbone for the other viruses and does not contain any inserted sequences in the duplicated subgenomic promotor region or the structural ORF. Arrows indicate subgenomic promoter transcription start sites. (B) MRE/rprORF and MRE/control contain sequences inserted after the final codon of the capsid sequence in the structural ORF. In both constructs, the first 9 nucleotides of PE3 are duplicated following the capsid to allow autoproteolytic activity of the capsid. Immediately following the PE3 insertion, MRE/rprORF contains a ubiquitin-*reaper* fusion with an HA tag. MRE/control contains ubiquitin and HA but does not contain *reaper*. The inserted sequence of both constructs ends with FDMV 2A which will release peptides as they are translated. (C) MRE/rpr contains the *reaper* gene inserted in the duplicated subgenomic promoter region. The figure was created with Biorender.com.

After these viruses were constructed, we first confirmed expression of the Reaper protein from MRE/rprORF by immunoblotting (Fig. 2). C6/36 cells were infected with MRE/rprORF, 5’dsMRE16ic (labeled MRE16 in the figure), or MRE/control, or they were mock infected, and at 24 or 48 hrs post-infection (hpi), protein was isolated and HA-tagged proteins were detected by immunoblotting. At 24 hpi we detected an HA-reactive protein in MRE/rprORF-infected cells of about 18 kDa. This likely represents uncleaved ubiquitin-Reaper fusion protein, which is predicted to be around this size. At 48 hpi, lysates from cells infected with MRE/rprORF still contained the 18 kDa protein but in addition had another protein of around 9 kDa, which likely corresponds to the HA-tagged Reaper protein after cleavage from ubiquitin. No proteins of similar size were detected in any of the other treatments. We also observed an HA-immunoreactive band of about 57 kDa at both time points in MRE/control-infected cells, as well as some faster migrating bands at 48 hpi (Fig. S1). Because in this control construct the HA tag is located immediately downstream of ubiquitin, we speculate that these bands may correspond to ubiquitin that was still fused to the HA tag and had been conjugated to unknown protein(s) by ubiquitin ligases. We did not observe any replication deficit in these constructs (see Fig. 4), suggesting that the expression of ubiquitin did not significantly affect virus replication. A protein of about 72 kDa that was seen in all treatments (Fig. S1) was assumed to be of cellular origin and detected as the result of antibody cross-reactivity.

**Fig. 2:**
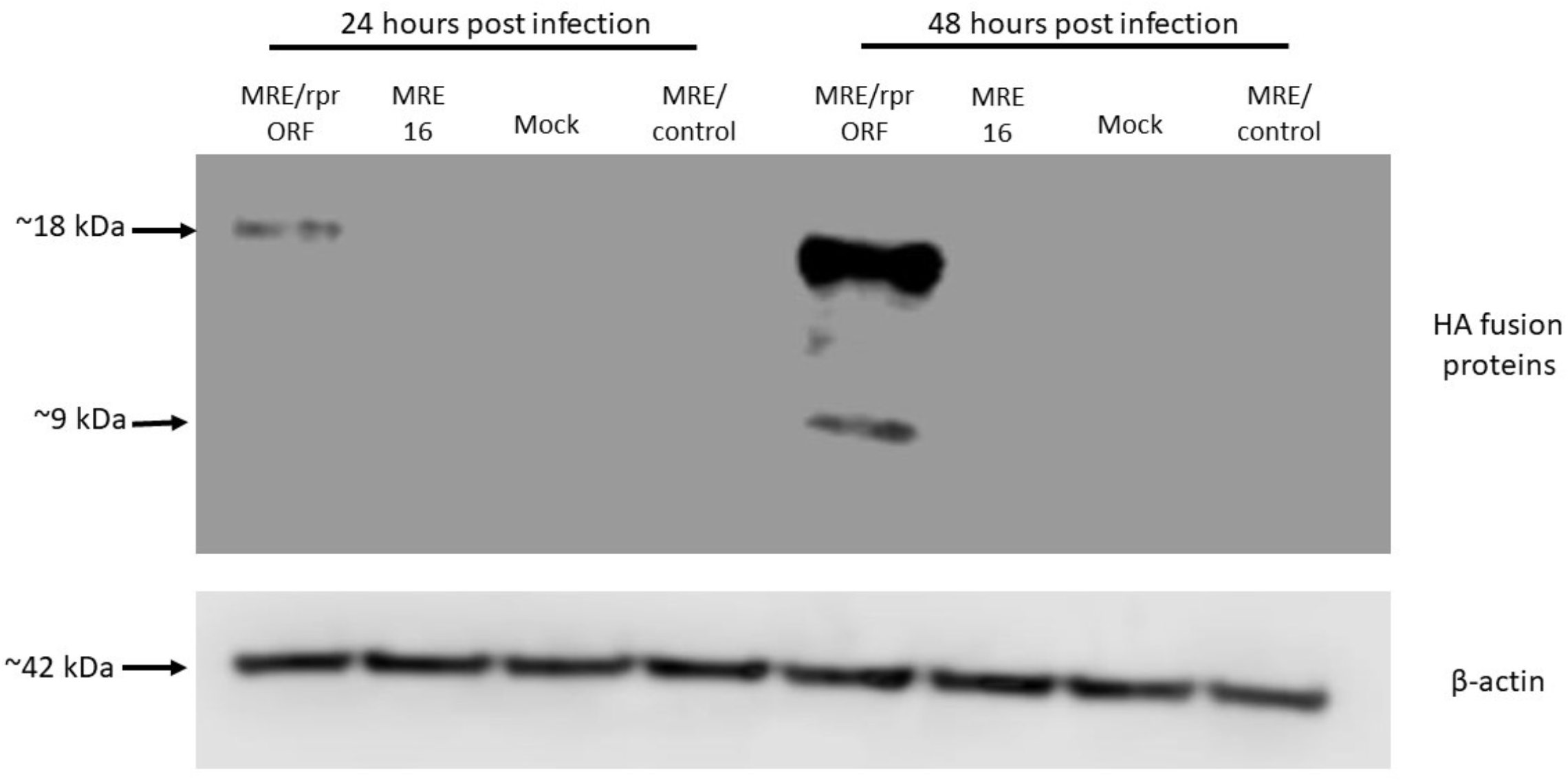
MRE/rprORF expresses the Reaper protein in infected cells. C6/36 cells were infected with MRE/rprORF, 5’dsMRE16ic (labeled MRE16), MRE/control or were mock infected and protein was extracted at 24 and 48 hpi. Immunoblotting was done using antibodies against HA (upper panel) or β -actin as a loading control (lower panel).

### MRE/rprORF causes increased apoptosis in C6/36 cells

To confirm that the Reaper protein expressed from MRE/rprORF was functional we infected cells with this virus and then tested for markers of apoptosis, as was previously done for MRE/rpr (22). We first demonstrated stimulation of apoptosis by MRE/rprORF infection using a chromatin fragmentation assay (Fig. 3A). Endonucleolytic chromatin fragmentation (or nucleosomal laddering), which is characterized by the appearance of a ladder-like pattern on gel electrophoresis, has long been known to be a hallmark of apoptotic cells (32–34). A nucleosomal ladder in multiples of approximately 140 bp was evident at 48 hpi in cells that were infected with MRE/rprORF, while DNA from cells that were infected with MRE/control or 5’dsMRE16ic (abbreviated as MRE16) did not display this pattern. We also demonstrated increased apoptosis in these cells using a caspase assay which measures effector caspase activity by detecting increased cleavage of a fluorogenic substrate, Ac-DEVD-AFC (Fig. 3B). We did not find differences in caspase activity between cells infected with MRE/rprORF, MRE/control or MRE/rpr when they were collected at 24 hpi. However, at 48 hpi, cells infected with MRE/rprORF or MRE/rpr were found to have increased effector caspase activity when compared to MRE/control. This increase between 24 and 48 hpi corresponds to the levels of Reaper expression in MRE/rprORF-infected cells as determined by immunoblotting (Fig. 2).

**Fig. 3:**
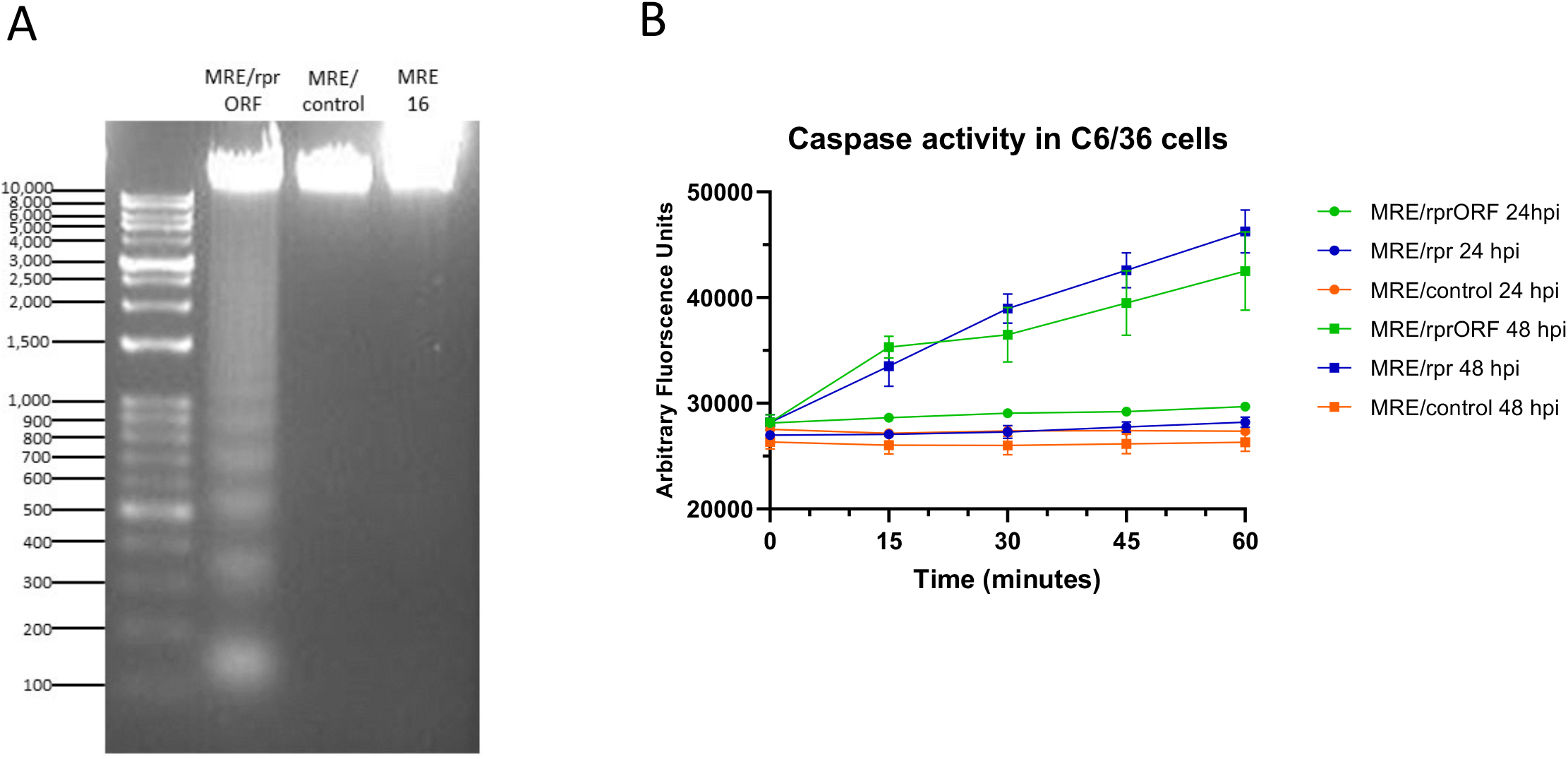
MRE/rprORF causes apoptosis in C6/36 cells. (A) C6/36 cells were infected with MRE/rprORF, MRE/control or 5’dsMRE16ic (labeled MRE16). After 48 hrs the cells were collected, DNA was extracted and run on an agarose gel containing ethidium bromide. (B) Cells infected with MRE/rprORF, MRE/rpr or MRE/control were collected and lysed at 24 and 48 hpi. Caspase activity from these cell lysates was measured by examining their ability to cleave the fluorogenic substrate Ac-DEVD-AFC. Fluorescence measurements were taken every 15 min for 1 hr. Three biological replicates were performed. Error bars indicate SEM.

**Fig. 4:**
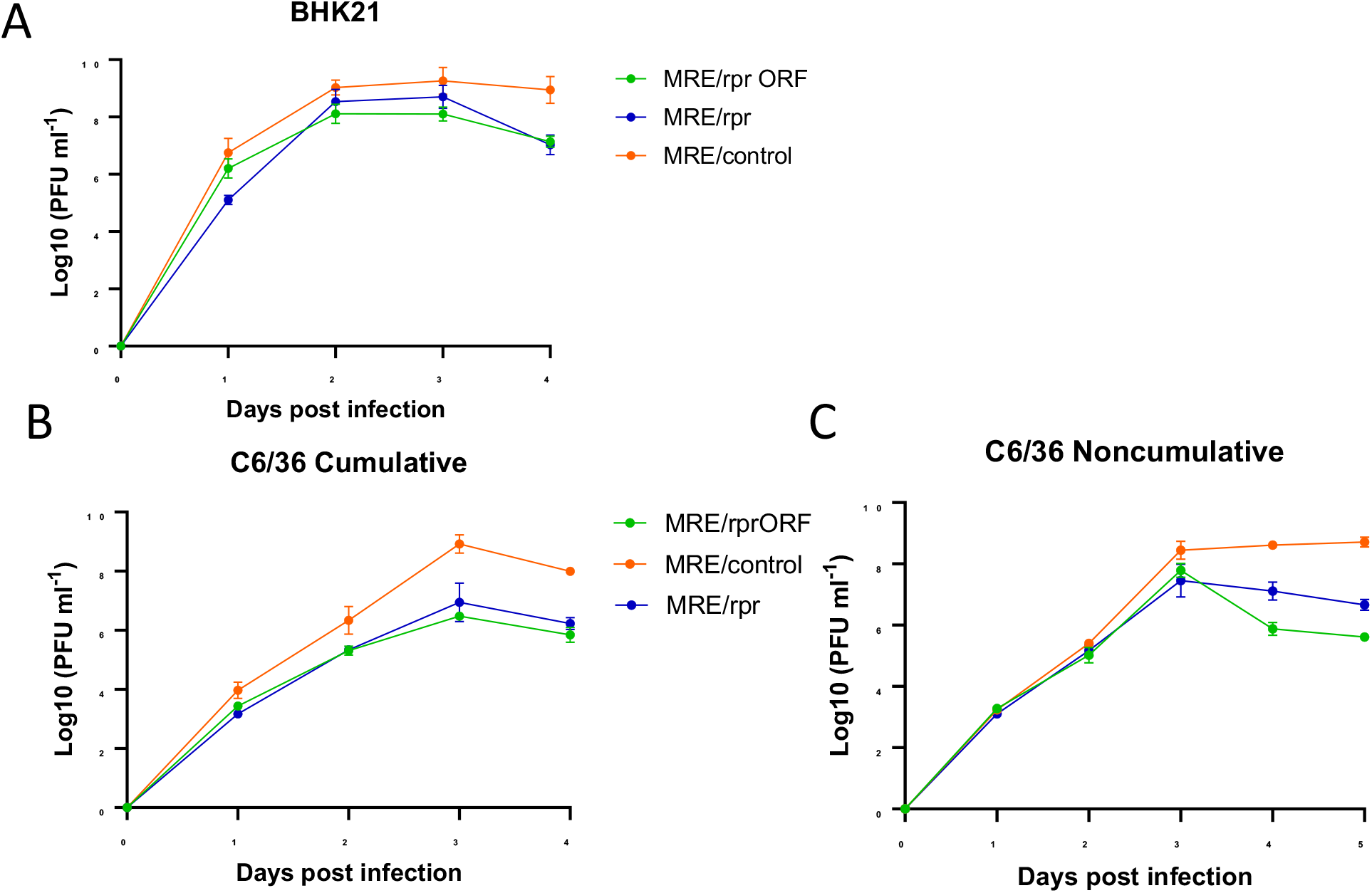
Virus growth curves in BHK-21 and C6/36 cells. (A-B) BHK-21 or C6/36 cells were infected with MRE/rprORF, MRE/rpr or MRE/control at an MOI of 0.1 and a sample of supernatant was removed and titered by TCID_50_ assay at each of the indicated timepoints. (A) In BHK-21 cells, MRE/rprORF (*p*=0.003) and MRE/rpr (*p*=0.0002) were significantly different compared to MRE/control. MRE/rpr and MRE/rprORF were not significantly different (*p*=0.9879). (B) In C6/36 cells, MRE/rprORF (*p*<0.0001) and MRE/rpr (*p*<0.0001) were significantly different compared to MRE/control. MRE/rprORF and MRE/rpr were not significantly different (*p*=0.7702). (C) Non-cumulative growth curve. C6/36 cells were infected with MRE/rprORF, MRE/rpr or MRE/control at an MOI of 0.1. At each indicated timepoint, a sample of supernatant was removed for TCID_50_ assay. The remaining cell culture medium was then removed from each well, the cells were rinsed, and the media was replaced. MRE/rprORF (*p*<0.0001) and MRE/rpr (*p*<0.0001) were significantly different compared to MRE/control. MRE/rprORF and MRE/rpr were also significantly different (*p*=0.0291). In A-C, three independent biological replicates were performed. Titers were log transformed and compared with 2-way ANOVA and Tukey’s multiple comparison test. Error bars represent SEM.

### MRE/rprORF and MRE/rpr show replication differences compared to control SINV in BHK-21 and C6/36 cells

To determine how Reaper expression would affect SINV replication and whether the insertion site of *reaper* in the viral genome would influence this effect, we infected BHK-21 and C6/36 cells and sampled the cell culture media by median tissue culture infectious dose (TCID_50_) assay at several timepoints to construct growth curves (Fig. 4). In BHK-21 cells we found that MRE/rprORF and MRE/rpr had reduced titers compared to MRE/control at most timepoints, with the largest difference being at 4 days post-infection (dpi) (Fig. 4A). There was no significant difference between the MRE/rprORF and MRE/rpr growth curves. In C6/36 cells, we constructed a cumulative growth curve as well as a non-cumulative growth curve in which the cells were rinsed after sampling at each timepoint (Fig. 4B). In the cumulative growth curve, both MRE/rprORF and MRE/rpr had lower titers than MRE/control at most timepoints and the differences were found to increase at later timepoints from about 10-fold at 2 dpi to around 100-fold at days 3 and 4. Differences between MRE/rprORF and MRE/rpr were not found to be significant. In the non-cumulative growth curve, MRE/rprORF and MRE/rpr titers were similar to MRE/control at early timepoints but the Reaper-expressing viruses diverged from MRE/control at the later timepoints of 4 and 5 dpi. Titers of MRE/control remained near 10^8^ PFU ml^-1^ after 3 dpi while MRE/rprORF and MRE/rpr titers reached between 10^7^ and 10^8^ PFU ml^-1^ at 3 dpi and then declined at 4 and 5 dpi. This decrease was expected since the Reaper-expressing viruses were causing apoptosis by these later time points. The growth curve of MRE/rprORF was found to be significantly different from MRE/rpr, with MRE/rprORF showing a more drastic decline after peaking at 3 dpi.

### Mosquitoes that ingest blood containing MRE/rprORF show increased caspase activity in midgut

We next wanted to see if we would find evidence of apoptosis in mosquitoes after they fed on blood containing the MRE/rprORF virus. In this experiment we allowed mosquitoes to feed on blood with MRE/rprORF, MRE/rpr or MRE/control. After three days the mosquitoes were dissected and pools of 8 midguts were tested for effector caspase activity. We found that there were significant differences in effector caspase activity between the three treatments by two-way ANOVA (Fig. 5). Pooled midguts from the mosquitoes that fed on blood containing MRE/rprORF had the highest effector caspase activity as indicated by the increase in fluorescence over time, suggesting that mosquitoes in this group had the most apoptosis occurring in their midguts. Caspase activity in MRE/rpr-exposed midguts was only marginally higher than in midguts exposed to MRE/control at this time point. Nonetheless, the results confirmed that MRE/rprORF infection induced apoptosis in mosquito midgut.

**Fig. 5:**
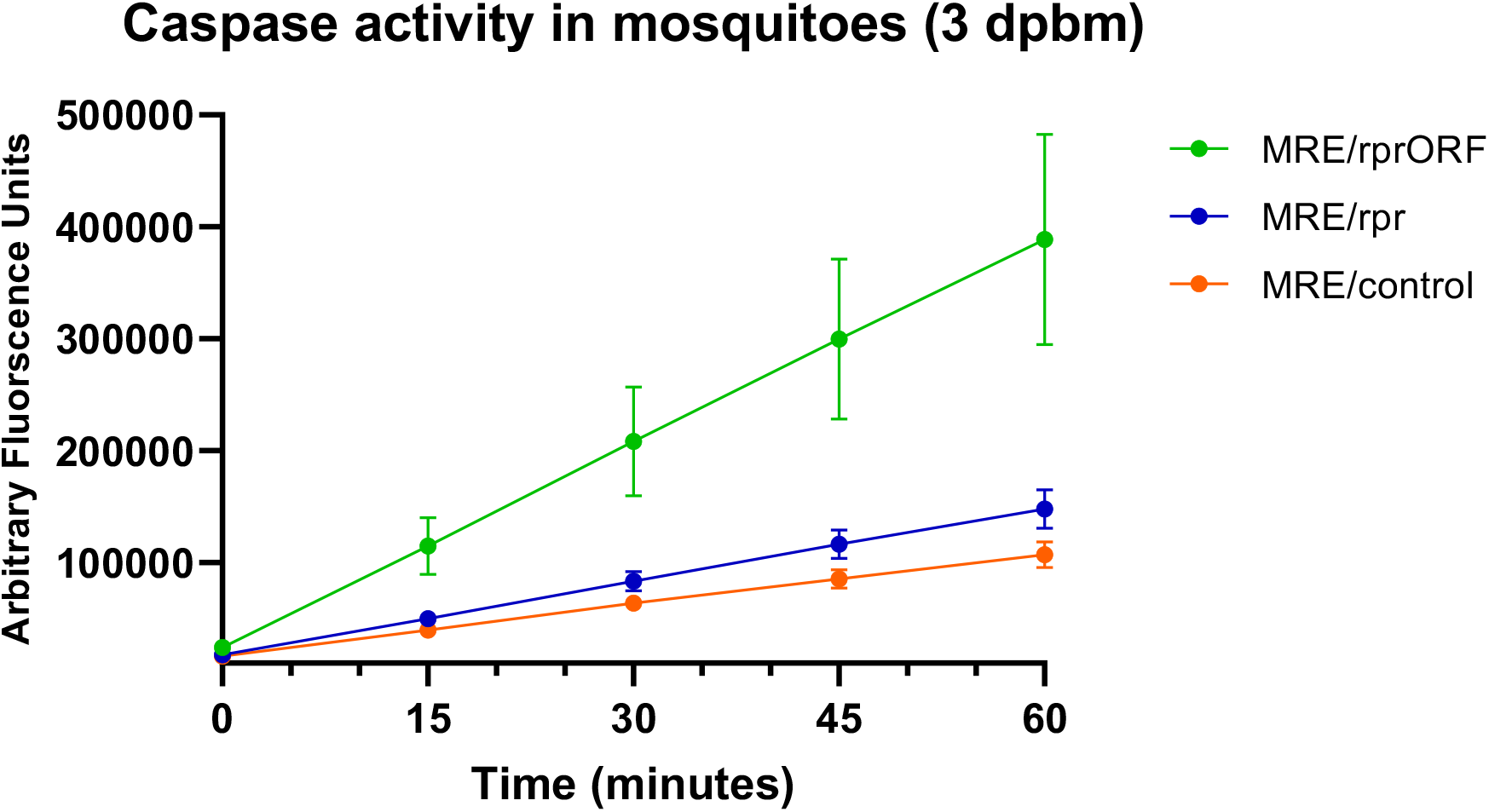
MRE/rprORF causes apoptosis in *Aedes aeygpti* midgut. Mosquitoes were fed blood containing MRE/rprORF, MRE/rpr or MRE/control and midguts were dissected after 3 days. Midguts were pooled in groups of 8 with 3 pools/treatment. Each pool thus represents an independent biological replicate. Cells were lysed by sonication and the supernatant was used for caspase assay. Cleavage of the Ac-DEVD-AFC substrate was monitored every 15 min for 1 hr. Viral treatment was judged to significantly contribute to variation using 2-way ANOVA (*p*=0.0055).

### MRE/rprORF is less able to infect mosquitoes, but replicates normally if infection is established

We then allowed mosquitoes to feed on blood containing MRE/rprORF, MRE/rpr or MRE/control for the purpose of comparing how many mosquitoes get infected and whether the infected mosquitoes show any differences in viral titer. To do this each mosquito midgut and carcass (defined as all remaining tissues of the mosquito other than the midgut), was titered by TCID_50_. In the midgut, we found that at all timepoints tested, the mosquitoes fed with MRE/rprORF were less likely to have detectable infection compared to MRE/rpr and MRE/control (Fig. 6A). This percentage increased over time in the MRE/rprORF-fed mosquitoes, from 25% of midguts with infection at 3 days PBM to 49% at 7 days PBM. By contrast, the percentage of midgut infection in the MRE/rpr-fed mosquitoes was not found to be significantly different compared to mosquitoes that fed on MRE/control, although it was lower at 3 days PBM (65% versus 81%). MRE/rpr infection also increased over time from 65% at 3 days PBM to 87% at 7 days PBM. Midgut infection percentage in the MRE/control-fed mosquitoes reached 81% by 3 days PBM and remained high over time.

**Fig. 6:**
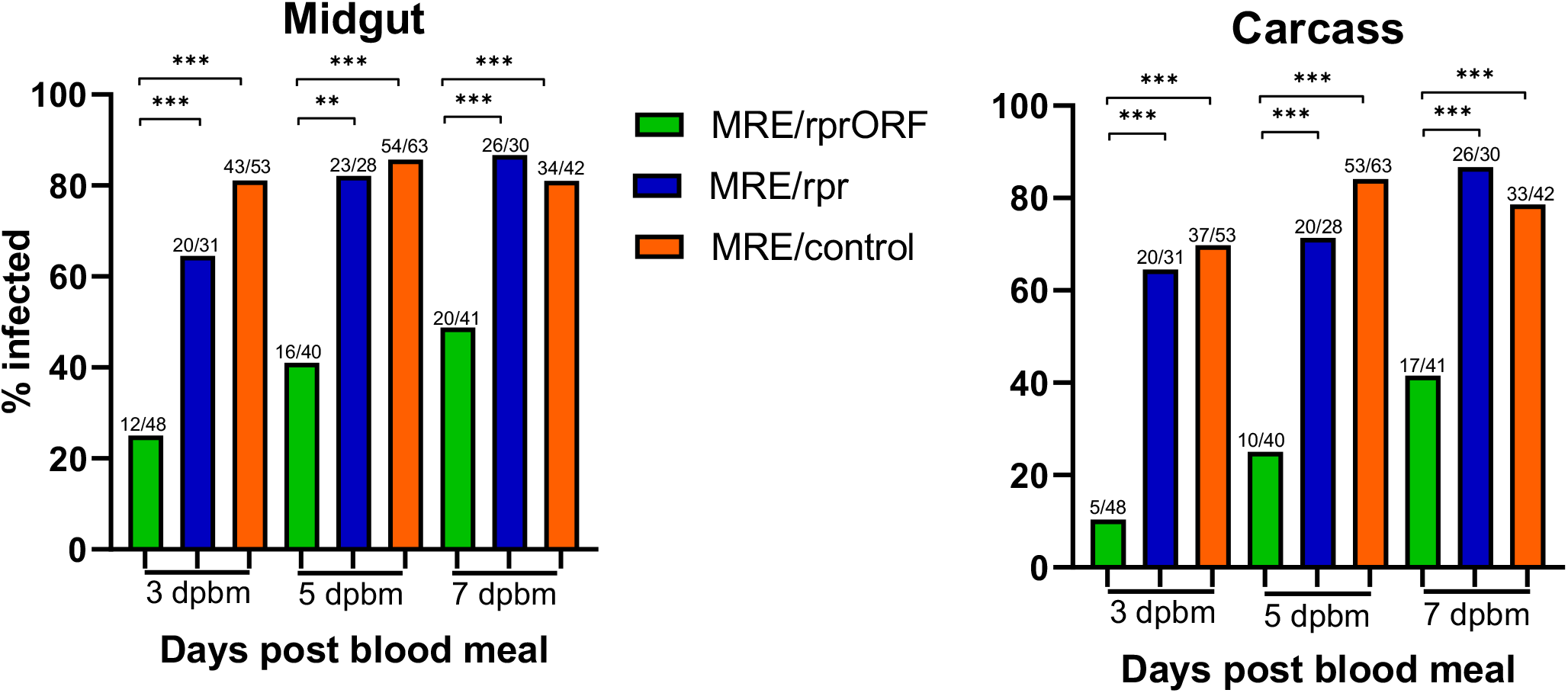
MRE/rprORF is less able to establish mosquito infection than MRE/rpr and MRE/control. Mosquitoes were fed blood containing MRE/rprORF, MRE/rpr or MRE/control and dissected at 3, 5 or 7 days PBM. Midguts (left) and carcasses (right) were titered by TCID_50_. Mosquitoes that did not have detectable titer were considered to be negative while any positive titer by TCID_50_ was considered to be positive. Treatments were compared using Fisher’s exact test (*** *p*<0.001, ** *p*<0.01). Statistical comparisons (brackets) are shown for all statistically significant differences.

Similar trends were seen in the percentage of mosquitoes with detectable carcass infections, representing dissemination of the virus from the midgut (Fig. 6B). The prevalence of mosquitoes with carcass infection in the MRE/rprORF-fed group was significantly lower compared to MRE/rpr and MRE/control at all timepoints tested. The percentage also increased over time and ranged from 10% at 3 days PBM to 41% at 7 days PBM. The prevalence of carcass infection in MRE/rpr-fed mosquitoes was not significantly different from MRE/control and ranged from about 65% at 3 days PBM to 87% at 7 days PBM. The percentage of carcass infection in MRE/control-fed mosquitoes was 74% at 3 days PBM with an increase to 84% at 5 days PBM and then a slight decrease to 79% at 7 days PBM.

To compare the titers in the midguts and carcasses between the three groups, we focused on only those mosquitoes that had detectable titers (Fig. 7). MRE/rprORF titers did not significantly differ from MRE/control at any of the timepoints tested in the midgut or carcass, while MRE/rpr titers were found to be higher than MRE/rprORF and MRE/control at 3 and 7 days PBM in the midgut and at 3 days PBM in the carcass. While statistical analysis of some of the titer data may be been affected by small numbers of infected mosquitoes, especially in the case of MRE/rprORF at the earlier time points, overall there did not appear to be any large differences in titer between the three viruses.

**Fig. 7:**
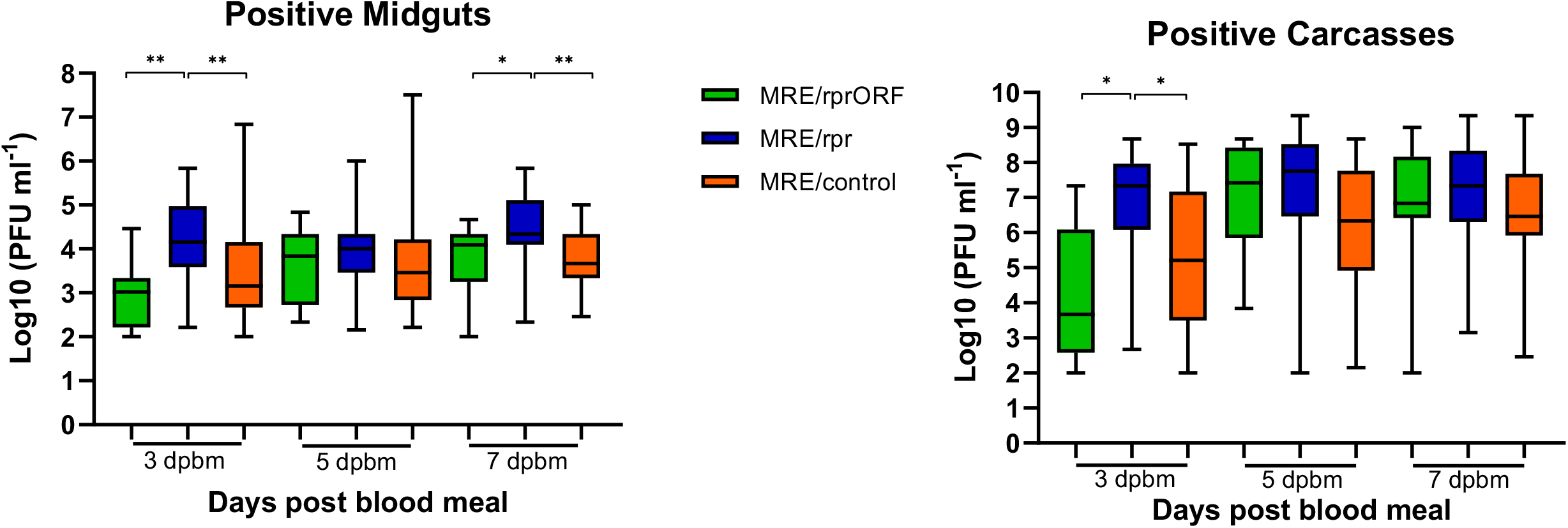
Mosquitoes infected with MRE/rprORF have similar titers compared to MRE/control. The titers of the infected midgut (left) and carcass (right) samples from Fig. 6 are shown. Titer values were log transformed and compared by 1-way ANOVA and Tukey’s multiple comparisons test. * *p*<0.05, \*\**p*<0.01, \*\*\**p*<0.001. Statistical comparisons (brackets) are shown for all statistically significant differences.

## DISCUSSION

The goal of this study was to generate a SINV construct that would more stably express the pro-apoptotic Reaper protein than the virus used in our previous studies and test how this virus would behave in cell culture and in mosquitoes. Previous work using a SINV construct that expressed Reaper from a duplicated subgenomic promoter (MRE/rpr) found that less mosquitoes developed disseminated infection than a control virus at early timepoints, but this difference disappeared over time due to deletions that accumulated in the inserted *reaper* sequence (18). Therefore, we were interested in learning what affect apoptosis would have on SINV infection if the Reaper protein was expressed in a more stable fashion.

The results of our study showed that expressing the Reaper protein from the structural ORF of SINV using a ubiquitin fusion strategy was successful. We were able to detect Reaper protein in MRE/rprORF-infected cells by immunoblotting, as well as increased markers of apoptosis in both infected cells and mosquito midgut. We detected what was likely the free Reaper protein at 48 hpi but also detected what was most likely the uncleaved ubiquitin-Reaper fusion protein at both 24 hpi and 48 hpi. Thus it appears that cleavage of ubiquitin fusion proteins in this system is relatively inefficient. We also detected a larger band of unknown origin in the MRE/control virus, which suggests that the ubiquitin expressed from these constructs may be utilized as a substrate by cellular ubiquitin ligases. We cannot conclude with certainty whether the expression of ubiquitin has any effect on virus structure or replication. However, the results of the growth curve experiments indicated that the ubiquitin fusion strategy did not result in any significant replication defects. In BHK-21 cells, MRE/control replicated to the highest level, with MRE/rprORF and MRE/rpr replicating to modestly lower levels before decreasing after 3 dpi. Although SINV naturally causes lytic cell death in BHK-21 cells, it is not surprising that Reaper expression would increase cell death to some degree, leading to lower titers. In the cumulative growth curve in C6/36 cells, the differences between MRE/control and the Reaper-expressing viruses were more pronounced, which was expected since SINV does not naturally cause a large amount of death in these cells (22). In this experiment the titers of MRE/rprORF and MRE/rpr closely matched each other, suggesting that the location of the *reaper* insert did not affect replication in these cells. In the C6/36 noncumulative growth curve, the titers of MRE/control, MRE/rprORF and MRE/rpr closely matched each other until 3 dpi, after which it appears that enough Reaper protein had accumulated to cause significant cell death and a decrease in titer was seen. MRE/rprORF decreased after 3 dpi to a greater extent than MRE/rpr, and these two growth curves were found to be significantly different by Tukey’s multiple comparisons test. The reason for this is not clear but it could be due to greater instability of the *reaper* insert in MRE/rpr, which may have decreased its effectiveness in causing cell death.

In mosquitoes, MRE/rprORF had a decreased ability to establish both midgut and carcass infection compared to MRE/rpr. At all timepoints tested, MRE/rprORF caused a lower infection prevalence compared to both MRE/control and MRE/rpr, while the prevalence of MRE/rpr infection did not significantly differ from MRE/control at any of the timepoints. This result differed from our previous study, which showed that the percentage of mosquitoes infected with MRE/rpr was lower than the control virus used in that study at early timepoints in both the midgut and the carcass (18). Although we observed that MRE/rpr infection prevalence was lower than MRE/control at 3 dpi (65% versus 81%), this difference did not reach statistical significance. Perhaps additional replicates would accentuate this difference. It is also possible that this difference could be due to different control viruses being used in the two studies. Regardless, the most important result for the purposes of this study was that MRE/rprORF showed less ability to establish infection in the midgut and disseminate compared to MRE/rpr. We conclude that this is likely due to the increased selective pressure to retain the insert in the structural ORF, since deletions within the *reaper* insert would have a 2 out of 3 chance of altering the reading frame and eliminating expression of the downstream envelope proteins, resulting in defective virus. In contrast, any MRE/rpr virus with a deletion within the *reaper* insert would remain viable.

The results of our study also differed from previous results when we examined the virus titers in mosquitoes that did become infected. We found that when MRE/rprORF was able to establish infection in the midgut, the titers were similar to MRE/control and this pattern continued in the carcass. One possible reason for this is that in the sub-population of mosquitoes that developed infection with MRE/rprORF, the *reaper* gene became non-functional due to mutations, despite the greater stability of the insert than in MRE/rpr. This possibility is supported by the increase in the percentage of mosquitoes that develop infection with MRE/rprORF over time, which suggests that as the virus replicates and is exposed to selective pressures, mutations may accumulate. Another possibility is that if the virus is able to accumulate to a certain level in midgut cells of some mosquitoes, apoptosis may not be able effectively limit viral replication and spread. When we examined the titers of mosquitoes that became successfully infected with MRE/rpr, we found that titers were higher compared to MRE/rprORF and MRE/control in the midguts at 3 and 7 days PBM and at 3 days PBM in the carcass. It is possible that in many of these MRE/rpr-infected mosquitoes the frequency of viruses with mutated *reaper* was higher, allowing it to replicate to higher levels compared to MRE/rprORF, but it is unclear why the titers would be higher than MRE/control. In any case, none of the three viruses appeared to have a large replication advantage over the others.

There were several limitations of our study which could be explored and improved upon in future studies. In this study we only looked at the effect of Reaper expression on SINV in *Aedes aegypti*. This should be explored in other virus/vector combinations to determine if the results of this study are generalizable to other situations. Additionally, while less than 50% of mosquitoes developed disseminated infection by 7 days PBM, we did see this percentage increase from 3 days PBM to 7 days PBM. It is possible that at further timepoints the percentage of infection of MRE/rprORF would become equivalent to control. The durability of this effect should be further explored in the future.

Overall, this study provides additional evidence that expression of the pro-apoptotic protein Reaper has a negative effect on the ability of SINV to establish infection in the midgut and disseminate to the carcass. Expressing Reaper by inserting the gene into the structural ORF caused a more robust reduction in infection prevalence than expression via the duplicated subgenomic promoter, with the difference in prevalence between MRE/rprORF and MRE/control being significant at all timepoints tested. These results provide additional evidence that the apoptotic pathway is antiviral in mosquitoes and possibly could be exploited to prevent transmission of arboviruses. Additionally, this study is consistent with previous findings that inserting genes into a viral ORF is a more successful strategy for durable expression of genes that are subject to negative selection.

## ETHICAL APPROVAL

This study did not involve human or vertebrate animal subjects.

## ACKNOWLEDGEMENTS

We thank Emma Francis, Abbey Phelps, Misty Bear and Jessica Rakijas for technical assistance. This is contribution 22-184-J from the Kansas Agricultural Experiment Station.

