## Supplemental Figure 1 for "Expressing the pro-apoptotic Reaper protein via insertion into the structural open reading frame of Sindbis virus reduces the ability to infect *Aedes aegypti* mosquitoes"

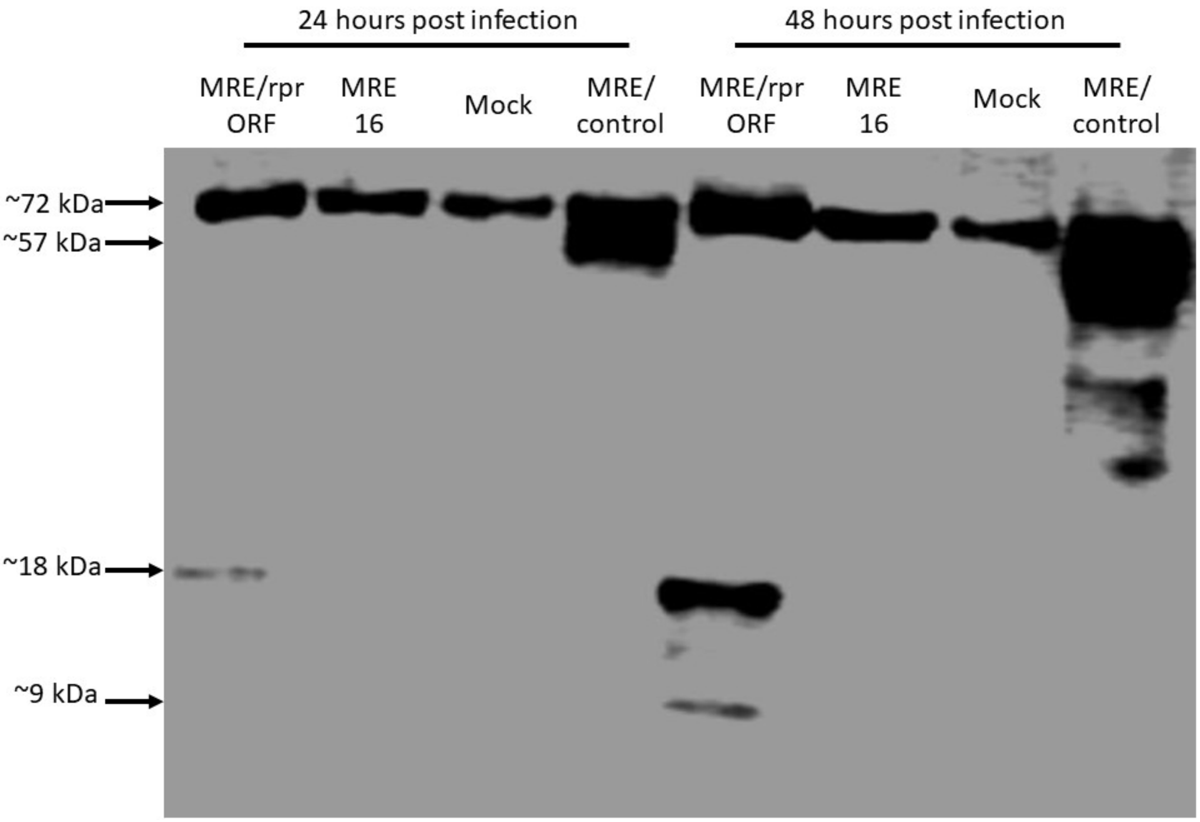


Fig. S1. Image of full immunoblot from Fig. 2. C6/36 cells were infected with MRE/rprORF, 5'dsMRE16ic (labelled MRE16), MRE/control or were mock infected and protein was extracted at 24 and 48 hpi. Immunoblotting was done using anti-HA antibody.
